# Multiple sources of signal amplification within the B cell Ras/MAPK pathway

**DOI:** 10.1101/415737

**Authors:** Justin D. Mclaurin, Orion D. Weiner

## Abstract

The Ras-Map kinase (MAPK) cascade underlies functional decisions in a wide range of cell types and organisms. In B cells, positive feedback-driven Ras activation is the proposed source of the digital (all-or-none) MAPK responses following antigen stimulation. However, an inability to measure endogenous Ras activity in living cells has hampered our ability to test this model directly. Here we leverage biosensors of endogenous Ras and ERK activity to revisit this question. We find that BCR ligation drives switch-like Ras activation and that lower BCR signaling output is required for the maintenance versus the initiation of Ras activation. Surprisingly, digital ERK responses persist in the absence of positive feedback-mediated Ras activation, and digital ERK is observed at a threshold level of Ras activation. These data suggest an independent analog-to-digital switch downstream of Ras activation, and reveals that multiple sources of signal amplification exist within the Ras-ERK module of the BCR pathway.

## Introduction

Digital or switch-like biochemical responses enable cells to convert gradual changes in external stimuli into binary cellular decisions such as cell cycle entry, differentiation, and programmed cell death (Spencer and Sorger, 2011; Tay et al., 2010; Huang et al., 2013). Positive feedback-driven protein activation is a common mechanism for generating digital signaling responses. Classic studies in Xenopus oocytes, for example, show how positive feedback within the MAPK cascade results in digital activation of the terminal kinase, p42 MAPK (Ferrell Jr. and Machleder, 1998). Subsequent studies have implicated digital MAPK responses in coordinating processes ranging from yeast mating responses to *Drosophila* tracheal placode invagination (Malleshaiah et al., 2010; Ogura et al., 2018). While the specific molecular details may vary across these organismal contexts, the result is the same: each makes use of positive feedback impinging on the MAPK machinery to drive switch-like like activation of the terminal kinase in the cascade (ERK in mammalian cells).

Cells of the B cell lineage also exhibit digital ERK activation, but here this binary response is thought to be generated by positive feedback at the level of Ras activation rather than within the MAPK cascade (Das et al., 2009). Here, the interplay of two Ras GEFs, RasGRP and SOS form the basis of a positive feedback loop to generate switch-like activation of Ras. Following antigen receptor triggering, RasGRP generates a small amount of initiating RasGTP, which is then amplified via RasGTP-driven activation of SOS (**Figure 1A**) (Margarit et al., 2003; Boykevisch et al., 2006). Computational models of this minimal Ras activating circuit posit a sharp transition (from off-to-on) in the Ras activation dose response curve, characteristic of switch-like activation (Das et al., 2009; Iversen et al., 2014; Jun et al., 2013). A potential result of this switch-like activation is a phenomenon known as hysteresis, a type of molecular memory, in which Ras activity persists in the absence of continuous antigen receptor engagement. However, live Ras activation dynamics in these cells has never been directly observed.

**Figure 1.**
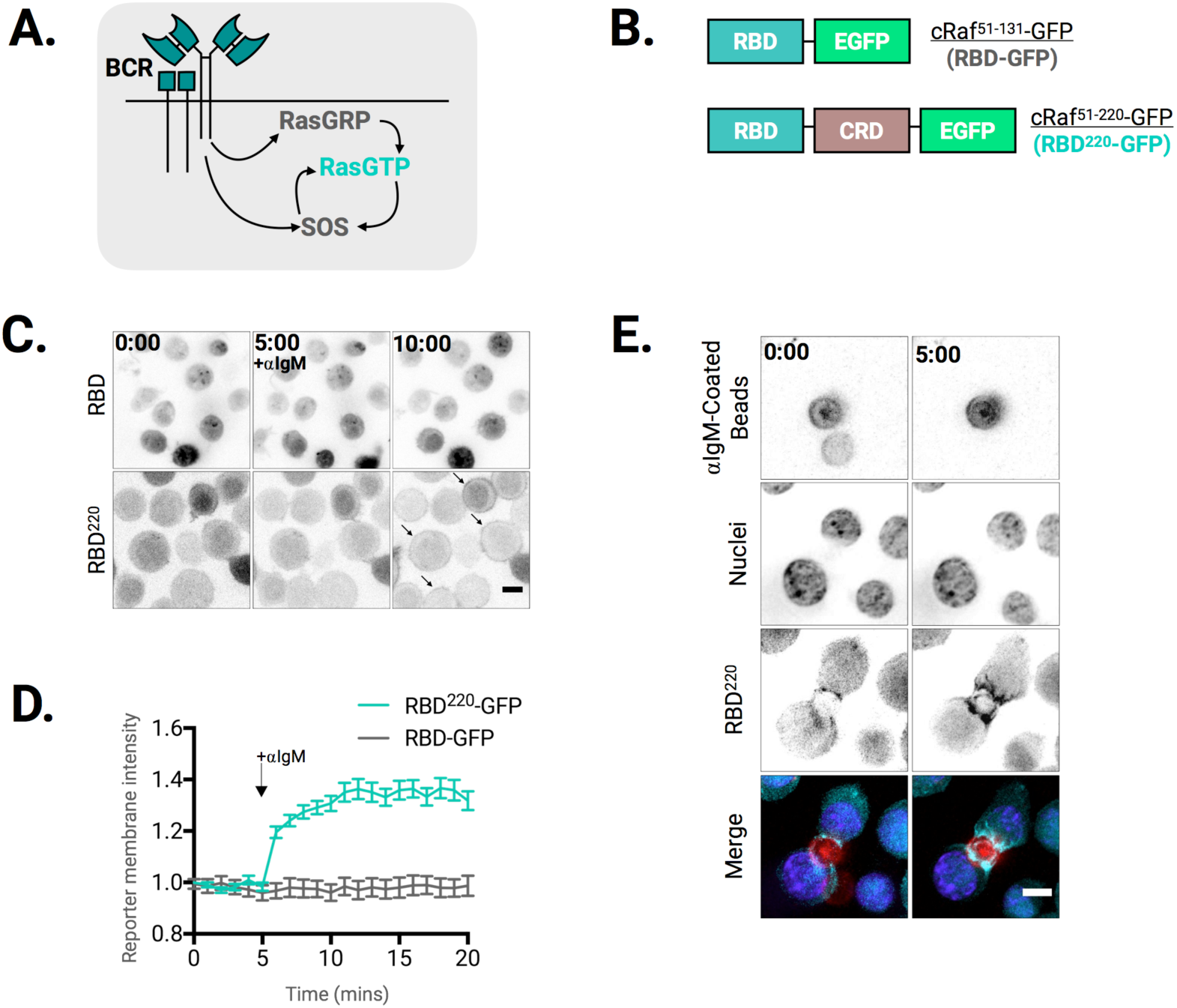
Visualizing Ras activation dynamics during B cell activation in living cells. **(A)** Schematic of the Ras GEFs that drive Ras activation downstream of B-Cell receptor triggering. **(B)** Domain structure of RasGTP biosensors. **(C)** and **(D)** Single cell image analysis of RasGTP biosensor localization following B cell activation. **(C)** Representative images of RBD-GFP (top) and RBD^220^-GFP (bottom) before and after addition of BCR agonist (αIgM, 10µg/mL). Scale bar is 5µm. Arrows highlight plasma membrane translocation of Ras biosensors. (**D)** Quantitation of normalized Ras biosensor intensity on the plasma membrane. Arrow indicates time αIgM addition (t = 5 mins). See supplemental methods for details on data collection and quantitation. **(E)** Representative images of RBD^220^-GFP localization in cells initially contacting BCR agonist αIgM-coated beads at the moment of contact (first column), and five minutes after contact (second column) Scale bar 5µm.

Here we take a live-imaging approach to analyze Ras-ERK signaling in individual Ramos B cells. We find that BCR engagement drives switch-like RasGTP responses at the single cell level, giving rise to bimodal Ras activation at the population level. Less receptor-based stimulation is required for the maintenance than for the initiation of a Ras response, providing evidence for hysteresis in Ras activation. Surprisingly we find that ERK responses remain binary even in the absence of positive feedback-driven Ras activation. This work supports multiple analog-to-digital switches in B cell activation, both at the level of Ras activation and between Ras activation and ERK activation.

## Results

### Visualizing Ras activity during B cell activation

Several groups have leveraged the high-affinity (∼20nM) interaction between the Raf-1 Ras binding domain (Raf1^51-131^, known as RBD) and RasGTP to generate FRET and membrane translocation-based reporters to quantify Ras activity in living cells (Chiu et al., 2002; Mochizuki et al., 2001; Oliveira and Yasuda, 2013). However, these approaches often lack the sensitivity to detect endogenous Ras and require overexpression of Ras proteins to produce detectable signal, potentially altering Ras regulation. Alternatively, endogenous RasGTP has been detected in live cells using a reporter in which RBD is multimerized (to increase avidity) and mutagenized (to decrease affinity), but this approach has the potential difficulty of responding to Ras density or sequestering endogenous Ras due to the high avidity of the multimeric reporter (Augsten et al., 2006; Rubio et al. 2010). To circumvent these issues, we make use of an extended fragment of cRaf/Raf1 that includes a second Ras binding site, the cysteine rich domain (CRD) (Thapar et al., 2004; Williams et al., 2000). This monovalent probe, Raf1^51-220^ (which we refer to as RBD^220^), was previously shown to detect endogenous levels of RasGTP in a variety of cell types with 1:1 stoichiometry (Anderson et al. 2011; Bondeva et al., 2002; Hibino et al., 2009). When expressed in Ramos B Cells, RBD^220^ tagged with eGFP (RBD^220^ -GFP) rapidly translocated to the plasma membrane following stimulation with BCR crosslinking F(ab’)_2_ fragments (αIgM) (**Figure 1B, C**). RBD-GFP, by contrast, failed to translocate to the membrane upon αIgM stimulation (**Figure 1B, C**). We observed an ∼37% increase in RBD^220^ membrane association following BCR crosslinking with saturating amounts (10μg/mL) of αIgM (**Figure1D**). GFP-tagged RBD, by contrast, showed no discernable membrane translocation or measurable cytoplasmic depletion to the same stimulus.

Previous work in which RBD^220^ was transfected into adherent cell lines suggested that this probe could inducibly, but possibly irreversibly associate with active Ras molecules on the membrane, thereby impairing downstream ERK activation (Bondeva et al., 2002). In contrast, for Ramos B cells stably expressing this probe we find that RBD^220^ membrane association is responsive to both the initiation and (as we will show in subsequent figures), the termination BCR signaling (**Figure 1D**). Furthermore, flow cytometry analysis of RBD^220^ expressing cells showed no alteration in the levels or kinetics of ERK phosphorylation compared to control cells, indicating that this Ras reporter does not perturb downstream signaling (**Figure S2A-C**) for the level of reporter expression used in our experiments. Pretreating Ramos cells with Syk (BAY-61-3606) inhibitor abolished αIgM induced RBD^220^ translocation to the membrane, while stimulating cells with the PKC/RasGRP agonist, PDBu, drove rapid reporter translocation (**Figure S3**). When RBD^220^ reporter cells were incubated with αIgM-coated beads, RBD^220^ localized to the region of contact between the cells and beads, reminiscent of an immunological synapse (**Figure 1E Supplemental Movie 2**). Together, these data indicate that RBD^220^ can be used as a sensitive dynamic reporter of BCR-stimulated Ras activation.

### Analog Ras activation driven by PKC/RasGRP

In B cells, Phospholipase C gamma 2 (PLCγ2) is recruited to the membrane following BCR ligation, where it catalyzes the cleavage of PI(4,5)P2 into Diacylglycerol (DAG) and Inositol trisphosphate (IP3). This DAG recruits and activates RasGRP and its activator Protein Kinase C (PKC) to initiate RasGTP production. Transfection studies performed with full-length RasGRP1 showed that overexpression of this protein produces linear increases in expression of the distal Ras-ERK signaling effector CD69 as a function of RasGRP1 expression level (Das et al., 2009). Similar experiments performed with the catalytic domain of SOS (SOS_cat_) showed that this protein induced exponential increases of CD69 expression as a function of SOS_cat_ expression level (Das et al, 2009.). While these and other experiments suggest that RasGRP proteins drive analog increases in RasGTP, the expression dynamics distal Ras-ERK effectors can serve as poor indicators for upstream Ras signaling dynamics (Wilson et al., 2017). Moreover, the activity of RasGRP proteins is subject to complex regulation by DAG, Ca^2+^ levels, and phosphorylation, making it difficult to infer how RasGTP responses are regulated by RasGRP activity alone (Iwig et al., 2013; Teixeira et al., 2003; Zheng et al., 2005).

To probe how Ras activation responds in the absence of SOS-driven feedback, we performed dose-response experiments with phorbol 12,13-dibutyrate (PDBu), a chemical mimetic of diacyl glycerol (DAG) that acts stimulates the Ras GEF RasGRP but not SOS (Lorenzo et al., 2000). Western blot analysis revealed titratable induction of RasGRP3 T133 phosphorylation (a PKC phosphorylation site) upon PDBu addition, validating this approach as a means of titrating RasGRP activity. Importantly, PDBu stimulation did not induce an increase in global phosphotyrosine or the recruitment of GRB2 to the membrane, both of which are prerequisites for SOS membrane recruitment and activation (**Figure S4A, B**). Moreover, pretreating cells with the PKC inhibitor Gö6983 abolished PDBu-driven RBD^220^ membrane recruitment and ERK phosphorylation, confirming the necessity of this enzyme in sustaining PDBu-driven RasGTP levels (**Figure S4C**).

To analyze how Ras signaling changes as a function of RasGRP activity at the single cell level, we stimulated RBD^220^ reporter Ramos cells with different concentrations of PDBu and quantified RBD^220^ cytoplasmic depletion via time lapse microscopy. These dose-response experiments showed an acute increase in RBD^220^ membrane recruitment that varied in mean amplitude as a function of PDBu dose (**Figure 2B, C**). While similar fractions of cells responded to all doses of PDBu (ranging from 83% at 0.01μM to 96% at 10μM) (**Figure S5B**), at the population level we observed a gradual increase in the distribution of mean RBD^220^ membrane intensity as the PDBu concentration increased (**Figure 2D**). Consistent with this, we found that all doses of PDBu drove unimodal increases in RBD^220^ membrane intensity (**Figure 2D**) as assessed by Hartigans’ dip test, a statistical test that distinguishes between bimodal and unimodal population distributions that has been previously used to distinguish between bimodal and unimodal signaling responses (Hartigan and Hartigan, 1985, Das et al., 2009; Jun et al., 2013). Fitting these data with a Hill function produced an estimated Hill coefficient (nH) of 0.5 (**Figure 2E**). These data suggest that PKC/RasGRP drive analog (non-ultrasensitive) RasGTP production when triggered in the absence of SOS, consistent with previous reports (Das et al, 2009).

**Figure 2.**
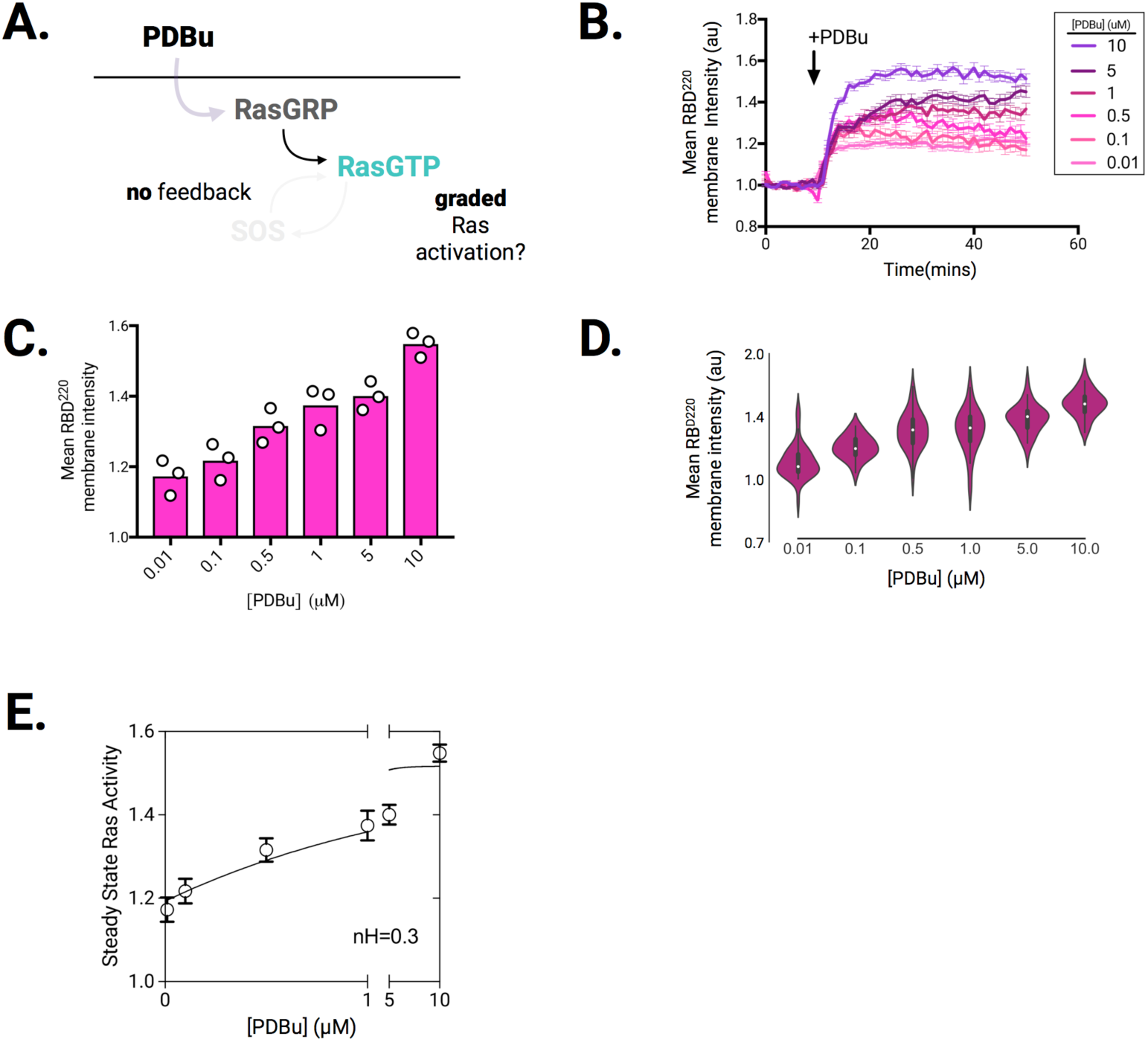
A water-soluble DAG mimetic (PDBu) drives graded, unimodal Ras activation responses. **(A)** Schematic of dose response experiment for cells exposed to step function increases of PDBu (RasGRP agonist) of different amplitudes. PDBu recruits RasGRP to the membrane and unlocks its catalytic activity. **(B-E)** Live cell analysis of RBD^220^-GFP membrane association (henceforth referred to as Ras activation) in response to PDBu. **(B)** Dose Response. Median intensity traces of cells stimulated with indicated doses of PDBu (see inset) over time. Arrowhead indicates time of PDBu addition (at ∼t=10mins). Error bars are SEM. Median intensity traces are generated from at least 50 cells per PDBu dose. Traces are representative of three independent experiments. **(C)** Mean RBD^220^ membrane intensity (calculated from t = 20-40mins) from cells stimulated as indicated in (B). Each circle represents the mean response from an individual experiment (n=3). **(D)** Violin plots of cells stimulated as indicated in (B), showing unimodal Ras activation responses at all PDBu doses. Mean response to indicated dose of PDBu (x-axis) was calculated as the mean membrane intensity on a per-cell basis between t=20-40mins (n=50 cells displayed per PDBu dose). **(E)** Steady-state Ras activity (see methods) across PDBu dose response experiments. nH (inset) indicates Hill coefficient, showing lack of Ras ultrasensitivity for PDBu input.

### Switch-like induction in BCR-driven Ras Responses

We next analyzed RasGTP signaling dynamics in the context of full BCR signaling. BCR ligation activates both RasGRP and SOS, and we reasoned that titrating BCR output could serve as a means of tuning the activity of these proteins. αIgM stimulation drove a dose-dependent increases in global phosphotyrosine production, validating our ability to titrate receptor output with this approach (**Figure 3A, S5A**). Low-level (0.25μg/mL and below) stimulation with BCR agonist elicited dose-dependent increases in the rate and mean amplitude of Ras activity, while high level (0.5μg/mL and above) stimulation yielded saturating response rates and amplitudes (**Figure 3B, C**). In addition, Ras responses were, on average, persistent (>50 minutes) at all αIgM doses applied.

**Figure 3.**
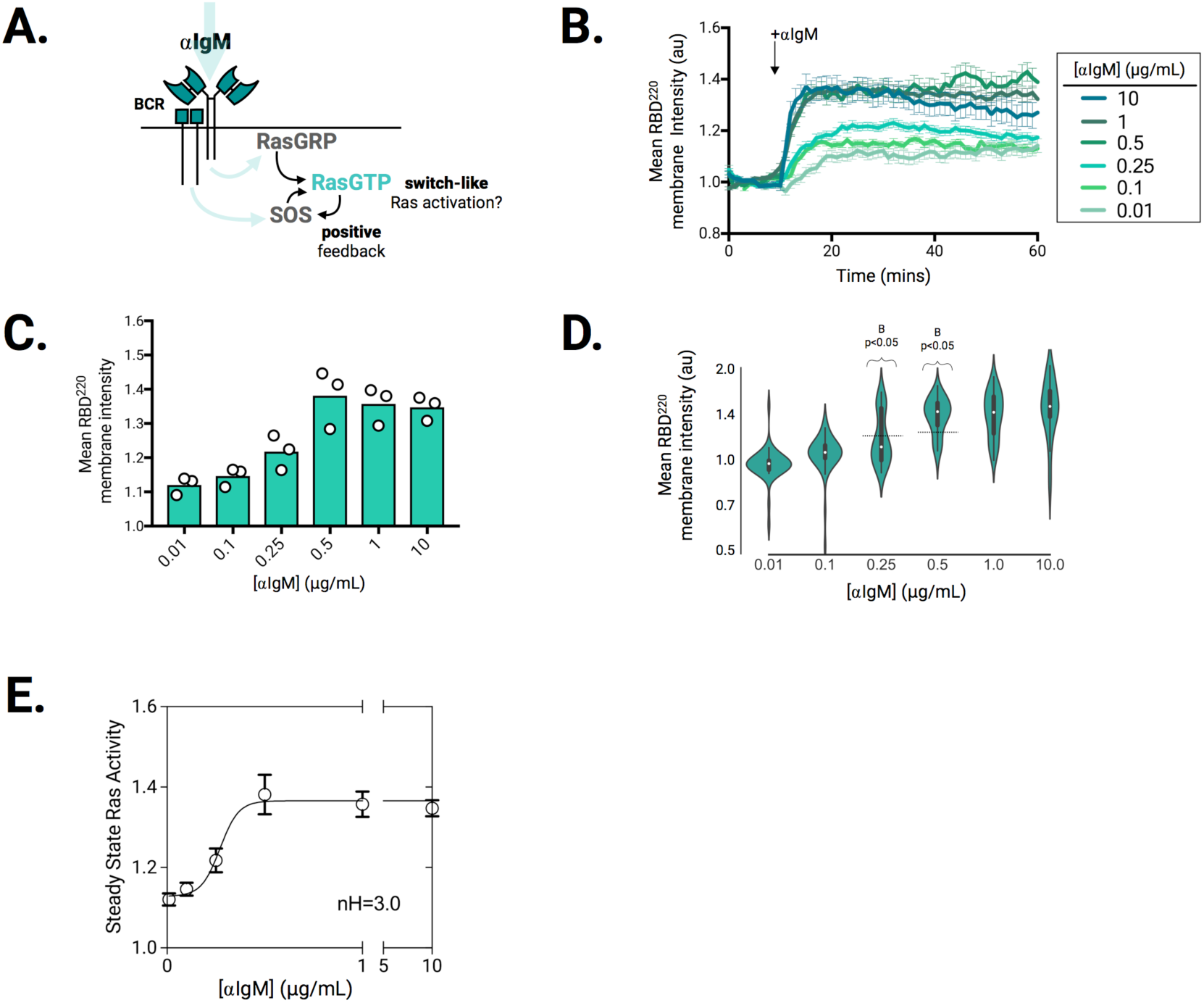
BCR agonist drives switch-like, bimodal Ras activation. **(A)** Schematic of dose-response experiment for cells exposed to step increases of BCR agonist (αIgM) of different amplitudes. **(B-E)** Live cell analysis of RBD^220^-GFP membrane association. **(B)** Mean intensity traces of cells stimulated with indicated doses of αIgM (see inset) over time. Arrowhead indicates time of αIgM addition (at ∼t=10mins). Error bars are SEM. Mean intensity traces are generated from at least 50 cells per αIgM dose. Traces are representative of three independent experiments. **(C)** Mean RBD^220^ membrane intensity (t = 20-40mins) from cells stimulated as indicated in (B). Each circle represents the mean response from an individual experiment (n=3). Violin plots of cells stimulated as indicated in (B) showing bimodal Ras activation at intermediate doses of αIgM. Steady-state response to indicated dose of αIgM (x-axis) calculated as the mean membrane intensity between t=20-40 mins (n=50 cells displayed per αIgM dose). Bimodality (B) and p-values were computed by Hartigans’ dip test (see methods). Dashed line indicates the dip statistic of cells stimulated with 0.25μg/mL and 0.5μg/mL αIgM (see methods). **(E)** Steady-state Ras activity (see methods) across αIgM dose response experiments. nH (inset) indicates Hill coefficient, demonstrating ultrasensitive Ras response for BCR agonist.

At the population level, the fraction of cells responding to αIgM stimulation ranged from 35% at 0.01μg/mL to 92% at 10μg/mL, increasing as a function of αIgM dose (**Figure S5D**). Hartigans’ dip test analysis level showed unimodal steady state Ras responses at the two lowest (0.01μg/mL and 0.1μg/mL) and highest (1μg/mL and 10μg/mL) doses of αIgM stimulation but bimodal distributions at intermediate doses (0.25μg/mL and 0.5 μg/mL, p<0.05, respectively) (**Figure 3D**). Consistent with these observations, steady-state Ras responses showed a sharp transition with an estimated Hill coefficient (nH) of 3, consistent with switch-like activation of Ras following BCR triggering (**Figure 3E**).

### Hysteresis in BCR-driven Ras activation

SOS-driven positive feedback is predicted to transiently maintain RasGTP levels (i.e. hysteresis) in cells in which signal flux through the BCR pathway is halted via antigen removal (Das et al, 2009.; Chakraborty et al., 2009). This hysteresis in Ras activity is presumably mitigated via stable association of SOS with the plasma membrane, as has been observed in several cell types (Christensen et al, 2016).

To date, most efforts have leveraged ensemble biochemical and fixed-cell readouts to analyze hysteresis in RasGTP responses downstream of the BCR. While these approaches have powerfully demonstrated B cell’s ability to maintain high levels of RasGTP in absence of persistent receptor triggering, several lingering questions warrant a reassessment of this model via the analysis of individual living cells: 1. Does maintenance of Ras activity depend on the relative timing with which cells experience BCR receptor triggering? 2. How heterogenous are hysteretic Ras responses across a population of cells? We paired our live imaging set up with a previously-developed means for reversibly controlling antigen receptor triggering to answer these questions (Weiss et al., 1987). The lectin Concanavalin A binds and crosslinks the BCR and has been previously used as a surrogate BCR ligand. BCR crosslinking by ConA can be rapidly attenuated via the addition of the high affinity ConA-binder Methyl-D-mannopyranoside (αMM), providing a means to induce and revert BCR signaling (**Figure 4A**). Indeed, we find that ConA rapidly induces RBD^220^ membrane association, and addition of αMM terminates Ras activation (**Figure 4B**). Importantly, BCR surface expression is required for ConA-induced signaling, as ConA does not drive RBD^220^ membrane association in IgM-deficient cells despite retaining its ability to associate with the cell surface (**Figure S6B**). These experiments validate ConA/αMM system as a reversible receptor triggering system compatible with our live imaging system.

**Figure 4.**
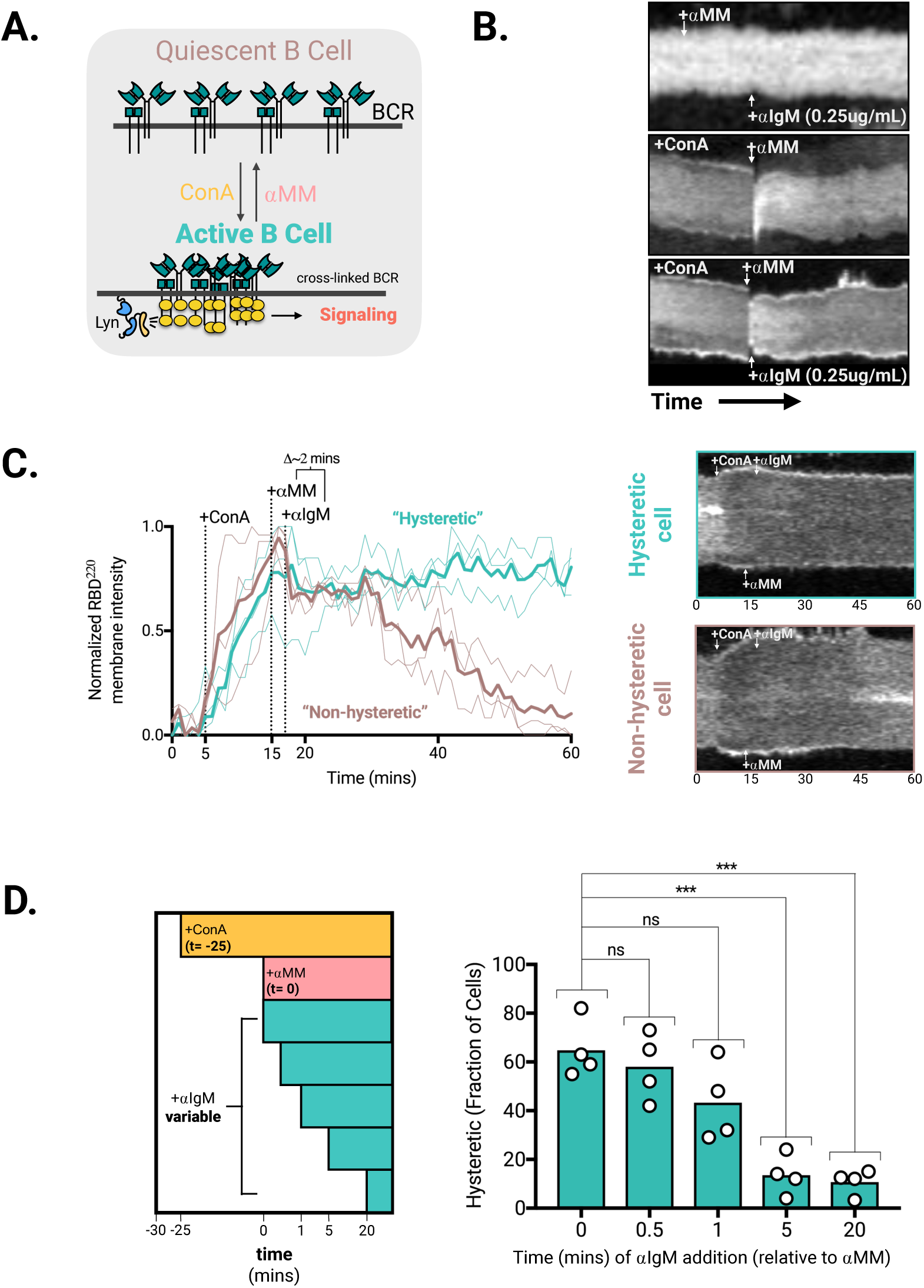
Hysteresis in BCR-driven Ras Activity. **(A)** Schematic of the reversible BCR triggering assay relying on ConA-mediated receptor crosslinking, and αMM-mediated reversal of B-cell receptor crosslinking. Lyn (a Src-family kinase) phosphorylates crosslinked receptors, leading to downstream signaling. **(B)** Single cell kymographs of RBD^220^-GFP over time. (**Top**) Cells stimulated with 100mM αMM (at t=2 mins), followed by 0.25μg/mL αIgM (at t=12mins). (**Middle**) Cells stimulated with 50μg/mL ConA (t=2mins) followed by 100mM αMM (t=12mins). (**Bottom**) Cell stimulated with 50μg/mL ConA (at t=2mins), 100mM αMM (t=12mins), and 0.25μg/mL αIgM (at t=13 mins). Experiment run length is 45 mins. **(C)** (Left) Single-cell traces showing examples of cells responding in a “hysteretic” (turquoise) or “non-hysteretic” (brown) manner following stimulation with 50μg/mL ConA (at t = 5 mins), 100mM αMM (t = 15 mins), and 0.25 μg/mL αIgM (at t=17 mins). (Right) Example kymographs of a hysteretic (top, turquoise border) and non-hysteretic (bottom, brown border) are displayed. **(D)** (Left) Schematic depicts relative time of addition of ConA (at t = 2 mins), αMM (at t = 27 mins), and αIgM (variable time points). (Right) Quantitation of the fraction of hysteretic cells as a function of time of αIgM addition relative to αMM addition. Hysteretic cells are defined as those whose Ras activity returns to within the 80^th^ percentile of their maximum post-ConA level following addition of αIgM. Each circle represents a single experiment. At least 20 cells were quantified per experiment. *** denotes statistical significance (p< 0.001, two-tailed unpaired student’s t-Test).

To determine whether less receptor signaling is required to maintain Ras activity than to initiate it, we crosslinked surface BCR with a high dose of ConA (50μg/mL), relieved those crosslinks with αMM (100mM), and re-crosslinked receptors with a low dose of αIgM (0.25 μg/mL) while monitoring RBD^220^ translocation dynamics. Importantly, we found that pretreating cells with 100mM αMM reduced the fraction of cells responding to saturating αIgM (10μg/mL) stimulus (**Figure S7A**), and 0.25 μg/mL αIgM was the minimum stimulus required to induce a Ras response in this context (**Figure S7B**). Ras activity was maintained in 59% of cells in response to iterative addition of ConA, αMM and αIgM (**Figure 4B-D**). We termed these cells “hysteretic” and defined them as those whose Ras activity returns to within the 80^th^ percentile of their maximum post-ConA level following the ConA/αMM/αIgM stimulus regimen (**Figure 4C**). Interestingly, we found that the delay between primary stimulus termination (αMM addition) and minimal stimulus application (αIgM) played a significant role in determining the fraction of hysteretic cells. Statistically similar, but, nevertheless declining fractions of hysteretic cells were found in instances where αIgM was added within ∼1 minute of αMM addition (**Figure 4D**). In contrast, addition of αIgM at 5 and 20 minutes post-αMM addition drove a statistically significant decrease in the fraction of hysteretic cells (**Figure 4D**), suggesting a temporal window in which Ras activity may be maintained before committing to decay.

### Signal Amplification downstream of Ras

How are switch-like Ras signals decoded by the downstream MapK machinery (**Figure 5A**)? SOS-mediated feedback is thought to drive the bimodal pattern of ERK phosphorylation seen in active lymphocytes (Das et al, 2009). Consistent with this, we found that Ramos B cells exhibit a bimodal pattern of ERK phosphorylation following BCR crosslinking with αIgM (**Figure 5B, left**). To test whether this binarization of ERK responses originates at the level of Ras activation, we supplemented our cell line co-expressing RBD^220^ with the recently described live cell ERK kinase translocation reporter (ERK-KTR) (Regot et al., 2014). Co-expression of these reporters in the same cell allowed us to track Ras and ERK signal activity with high temporal precision within individual cells (**Figure 5C**, **Supplemental Movie 3**). Time-course imaging experiments revealed a mean lag time of ∼4 mins between peak RBD^220^ membrane association and peak ERK-KTR activity in cells stimulated with a saturating dose (10 μg/mL) of αIgM (**Figure S8A**). Low doses of αIgM (0.01μg/mL and 0.1μg/mL) rarely increased ERK-KTR response despite driving detectable increases in RBD^220.^ membrane association (**Figure 5D**). Interestingly, these low doses of αIgM stimulus generally drove a <20% increase in RBD^220^ membrane association, suggesting that a threshold increase in Ras activity above basal levels may be required to trigger an ERK response in these cells. Stimulating cells with a higher dose (1μg/mL) of αIgM by contrast drove a >20% increase in RBD^220.^ membrane association and also induced potent ERK-KTR responses (**Figure 5D, bottom right**). At the single cell level, we found that greater than 20% increases in Ras activity corresponded to induction of an ERK-KTR response (**Figure 5E**). Kernel density estimation (bandwidth = 0.5) of these data showed that >20% increases in Ras activity corresponded >4-fold increase in median ERK kinase activity, with the population density exhibiting a bimodal transition (**Figure 5F**).

**Figure 5.**
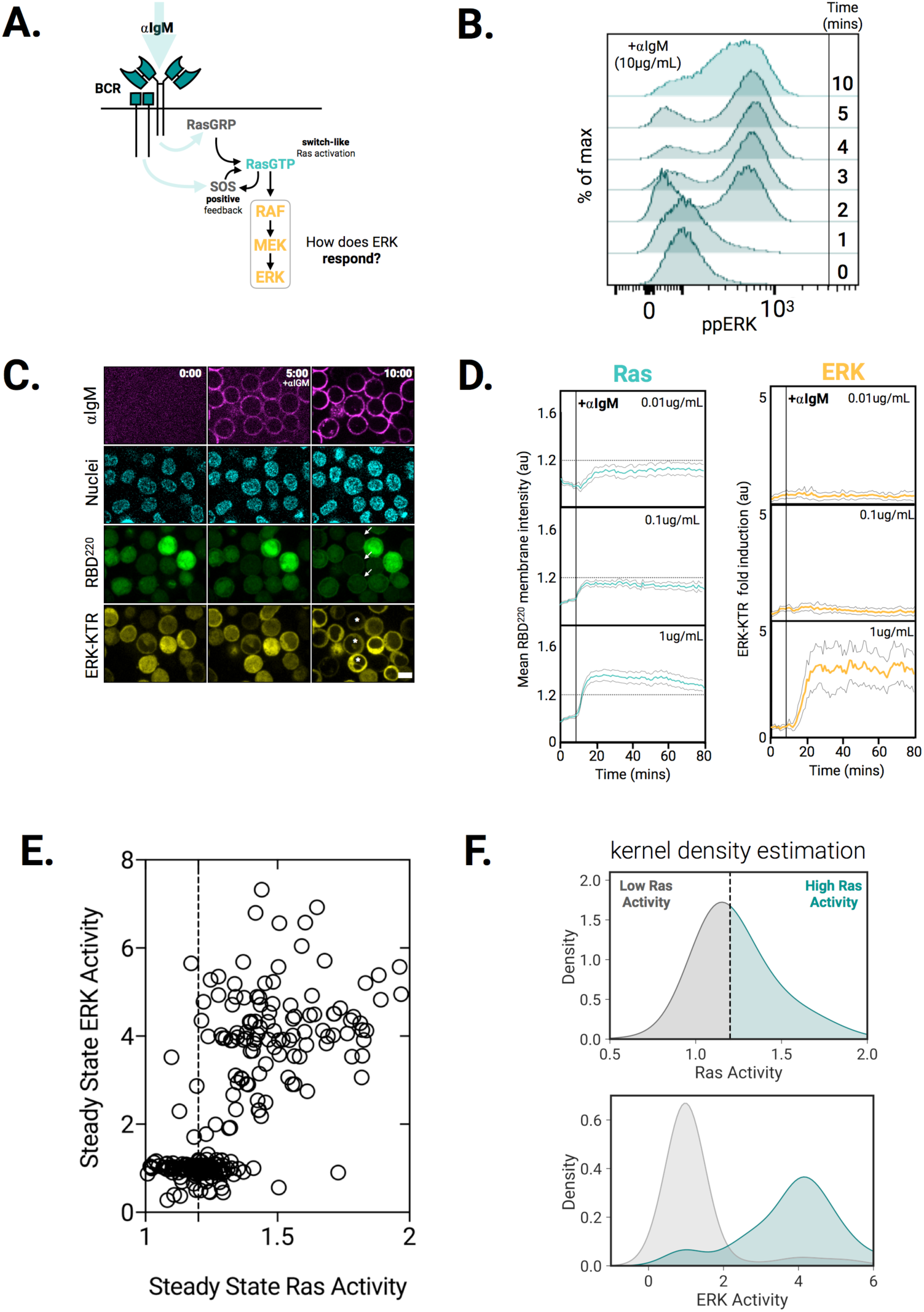
Bimodal ERK activation beyond a threshold level of Ras Activation. **(A)** Schematic depicting switch-like Ras activation arising from BCR-generated Ras positive feedback. **(B)** ERK phosphorylation time course from wildtype Ramos B cells stimulated with αIgM (10µg/mL, right) (representative of three independent experiments). **(C)** Representative (n = 2) time course images of αIgM (top, magenta), nuclei (second row, teal), RBD^220^-GFP (third row, green), and ERK-KTR-TagRFP-T (yellow, bottom) following APC-αIgM addition (10µg/mL). Nuclei are labeled with nucblue (see methods). 5µm scale bar. **(D)** Cells co-expressing RBD^220^-GFP (left, purple) and ERK-KTR-TagRFP-T (right, brown) were stimulated with indicated doses (inset) of αIgM. Mean response (50 cells per trace; pooled from three independent experiments) plotted over time. 95% confidence interval (black lines) envelope mean traces. Dashed lines (left column) indicate apparent threshold of Ras activity required to trigger ERK response (∼20% increase above basal). **(E)** Dose-response experiments. Representative single-cell mean steady-state ERK activity plotted as a function of mean steady-state Ras activity in individual cells. Each circle is data from an individual cell. Data are pooled from αIgM dose response experiments where cells were stimulated with 0.01, 0.1, 0.25, 1 or 10μg/mL αIgM. 50 cells per αIgM dose are plotted (250 cells total). Dashed line indicates 20% increase above basal Ras activity. **(F)** Kernel density estimations of Ras and ERK activity plotted in (E). Ras activity (top) is segmented into low and high by line (vertical, dashed) demarcating a 20% increase above basal activity. ERK activity (bottom) is plotted as a function of low (grey) and high (teal) Ras activity.

Bimodal responses are generated by switch-like systems responding in an all-or-none manner to stimuli above a threshold, while ignoring sub-threshold stimuli (Ferrell Jr. and Ha, 2014). While our analysis would be consistent with switch-like activation of Ras simply propagated to ERK without further amplification, it is formally possible that bimodal ERK responses are generated downstream of Ras via independent feedback modules (Bhalla et al., 2002; Ferrell Jr. and Machleder, 1998; Shin et al., 2009; Shindo et al., 2016). To test this idea, we stimulated cells with PDBu (a condition in which Ras positive feedback is not engaged and for which Ras activation is graded, not bimodal) and analyzed the resulting ERK responses in single cells (**Figure 6A**). Surprisingly, we found that in PDBu-stimulated Ramos cells, bimodal ERK phosphorylation is retained and is kinetically similar to BCR-activated cells (**Figure 6B, 5B**). Importantly, we found that this bimodal activation is preserved across a wide range of PDBu doses in both Ramos and Jurkat T cells, suggesting conservation of the underlying signaling network architecture leading to bimodal ERK downstream of unimodal Ras (**Figure S8B right panel, C**). This led us to hypothesize that ERK may respond in a digital manner to changes in Ras activity. To test this, we performed two-step experiments in which we pulsed cells sequentially with low (0.1μM) and high (1μM) doses of PDBu, reasoning that analog increases in Ras activity driven by PDBu would allow for analysis of ERK responses in the absence of upstream feedback (**Figure 6C**). Live imaging of RBD^220^ showed two step-like increases of plasma membrane translocation in response to sequential pulses of low and high PDBu stimulus, consistent with analog Ras activation (**Figure 6D, 6E bottom**). In contrast (for the same cells), ERK-KTR responded to either low or high pulses of PDBu, with each cell exhibiting a single step-like response despite experiencing two-pulses of stimulus (**Figure 6D and 6E, top**). Taken together, these data suggest that an increase in Ras activity above a threshold is necessary to trigger downstream ERK signaling. Furthermore, the Ras/MAPK cascade contains two independent modules for generating bimodal responses-- one within and one downstream of Ras activation.

**Figure 6.**
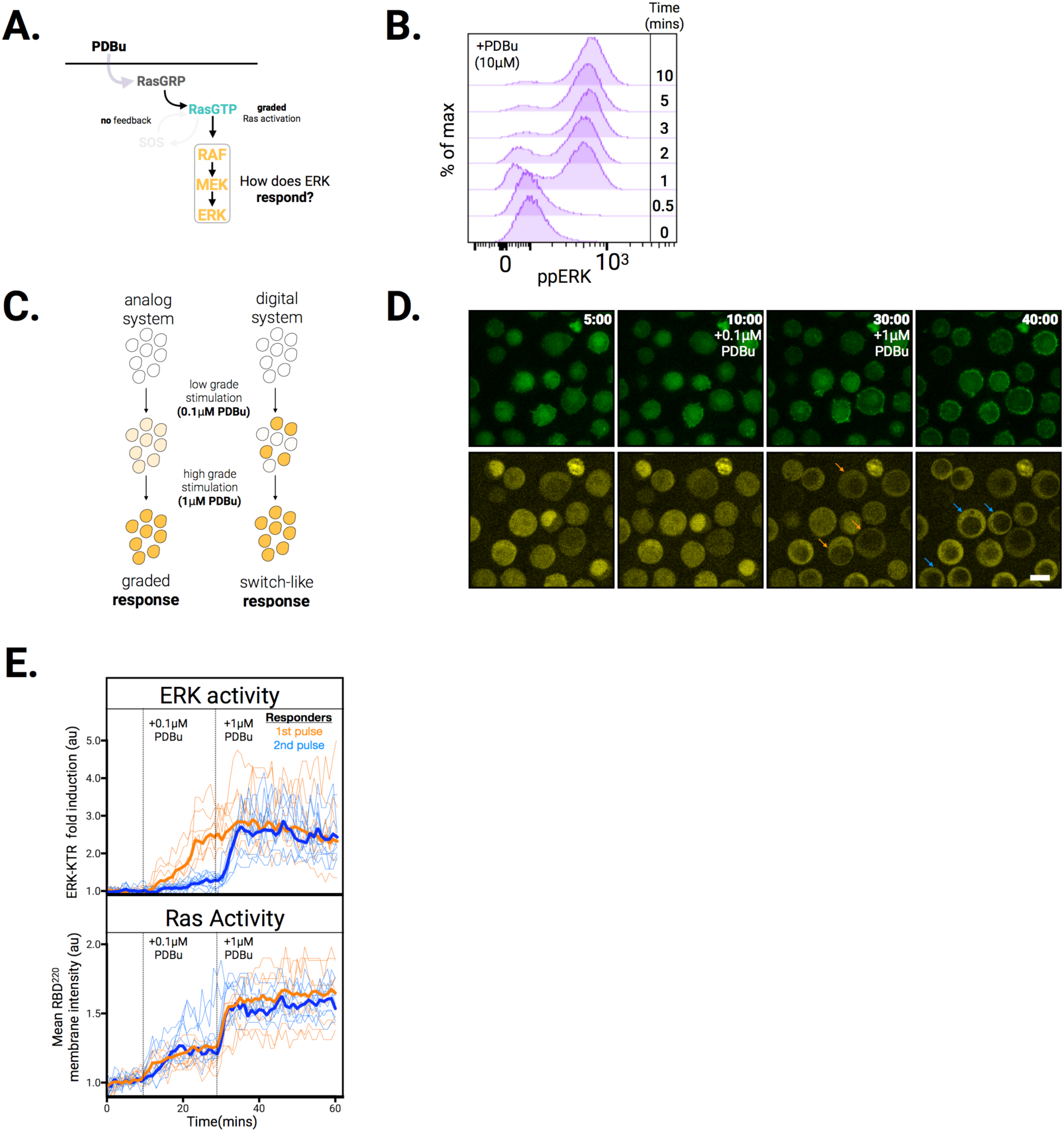
Digital ERK activation downstream of Ras. **(A)** Schematic depicting graded Ras activation driven by PDBu-mediated activation of RasGRP (this agonist does not engage the Ras positive feedback loop). **(B)** ERK phosphorylation time course from wildtype Ramos B Cells stimulated PDBu (10µM, right) (representative of three independent experiments). **(C)** Possible outcomes of an experiment in which analog (left) and digital (right) cellular systems are pulsed sequentially with low and high doses of PDBu. Graded increases are characterized by gradual increases in cell responses (left, yellow coloration). Switch-like responses are characterized by all-or-none responses on a per-cell basis (right, yellow coloration). **(D)** Representative (n=3) time course images of RBD^220^-eGFP and ERK-KTR-TagRFP-T Ramos cells stimulated with 0.1μM (at t=10mins) and 1μM PDBu (at t=30mins). Arrow heads indicate ERK-KTR expressing cells responding to the first (orange) or second (blue) pulse of PDBu. **(E)** Single cell traces of cells stimulated as in (D) depicting Ras (top) and ERK (bottom) responses. Traces are color coded to indicate cells whose ERK-KTR responses initiated at the first (orange) or second (blue) pulse of PDBu.

## Discussion

In this work we provide evidence for multiple signal amplification steps occurring at the level of Ras and ERK during B cell activation. BCR-driven Ras activation is highly switch-like (nH=3), while PDBu driven Ras activation is graded (nH=0.3), supporting the model that cooperation between RasGRP and SOS gives rise to nonlinear Ras pathway activation (Das et al, 2009.) **(Figure 2E, 3E)**. Less receptor-based stimulation is required to maintain a Ras response than to initiate it, consistent with hysteresis in BCR-RasGTP dose response curve (**Figure 3C** and Das et al, 2009). ERK activation is bimodal in BCR-triggered Ramos cells, and these bimodal responses proceed following Ras activation beyond a strict threshold **(Figure 5B, E)**. Surprisingly, ERK regulation is digital in its own right, exhibiting all-or-none activation even for inputs that fail to generate digital Ras responses **(Figure 6B, E)**. While these findings broadly support the predominant model for digital Ras activation in active lymphocytes, they raise important questions concerning the mechanistic basis for digitization of ERK signaling.

Feedback activation of SOS by RasGTP is the predominant mechanism invoked to explain bimodal nature of activation of Ras pathway effectors (e.g. ERK, CD69). However positive feedback and amplification at the level of Ras activation is just one mechanism capable of generating bimodal responses. Signaling protein oligomerization and phase separation, for example, provide additional mechanisms for initiating non-linear biochemical responses (Li, Banjade, Chen et al., 2009). Indeed, recent studies highlight the ability of GRB2 and SOS to form phase separated aggregates in activated T-Cells (Su et al., 2016; Huang et al., 2016). Moreover, active Ras molecules have been shown to form homotypic nanoclusters capable of digitizing downstream ERK signaling in EGF stimulated Baby Hamster Kidney cells (Tian et al., 2007). It will be interesting to evaluate how each of these mechanisms contribute to the Ras pathway signaling dynamics we describe here.

We previously used a set of optogenetic tools to characterize the signal transmission between active Ras and downstream ERK in 3T3 fibroblast cells (Toettcher et al., 2013). There we found near-linear transmission of Ras signals downstream to ERK nuclear localization. In contrast, our data here support a model in which an independent analog-to-digital converter drives ERK activation once active Ras exceeds a threshold level. What mechanisms might explain this discrepancy? In Xenopus, positive feedback within the MAPK cascade mediates digital pathway activity, while in yeast, a zero-order ultrasensitivity in the disassociation of Fus3 (the yeast MAPK) from the scaffold protein Ste5 mediates switch-like responses (Ferrell Jr. and Machleder, 1998, Malleshaiah et al., 2010). Pairing optogenetics with genomic and proteomic approaches will provide a framework for uncovering these differences in Ras pathway network architecture that underlie the differences in signaling responses between lymphocytes and fibroblasts (Toettcher et al., 2013; Wilson et al., 2017).

We have shown that analog Ras signals give rise to digital ERK responses, suggesting that consecutive digital switches exist at the level of Ras and ERK activation during normal BCR signaling. Why organize the Ras-MAPK pathway in this manner? One possibility is that switch-like Ras responses serve to digitize activation of other Ras effectors (besides Raf), such as PI3K and PKCζ. As bimodal Ras responses bracket the switch point for ERK kinase activity (**Figure 5B, E**), it is also possible that these sequential switches could confer robustness to MAPK activation. Lastly, lymphocytes are thought to integrate antigen signals in time during serial APC engagement (Zikherman and Au-Yeung. 2015). Hysteresis in Ras activity could act as a ‘molecular memory’ of previous antigen exposure. Our work provides a foundation for investigating how these multiple sources of signal amplification enable B cells to convert the magnitude, duration and frequency of antigen exposure into appropriate cellular activation.

## Materials and Methods

### Cell Culture

Ramos cells were obtained from the American Type Culture Center (ATCC). Jurkat cells were a kind gift from Art Weiss (UCSF). Both Jurkat and Ramos cells were maintained in 10G-RPMI (RPMI 1640 supplemented with L-Glutamine and 25 mM HEPES (Mediatech) and containing 1% (vol/vol) GlutaMAX (Gibco) and 10% (vol/vol) Fetal Bovine Serum (Gibco)). Cultures were maintained in a 37°C/5% CO_2_ incubator at a density ranging from 0.2-1 million cells/mL. HEK-293T cells (used to generate lentivirus for transduction) were maintained in DMEM (Mediatech) supplemented with 10% (vol/vol) Fetal Bovine Serum (Gibco). Cells were imaged in imaging media (RPMI 1640 without phenol red supplemented with 25mM HEPES, 1% L-glutamine and 0.1% fetal bovine serum).

### Plasmids

All constructs were cloned into the pHR lentiviral backbone (kindly provided by Ron D. Vale, UCSF containing the spleen focus forming virus (SFFV) promoter via standard Gibson assembly. Human cRaf/Raf1 cDNA was a gift from William Loomis (UCSD). RBD^51-131^ and RBD^51-220^ coding sequences were amplified from this cDNA and subcloned into a GFP containing pHR vector to generate pHR-RBD-GFP and pHR-RBD^220^-GFP plasmids, respectively. The ERK^KTR^ sequence was amplified from pLentiCMV-Puro-ERK^KTR^-Clover (Addgene # 59150) and subcloned into a pHR vector containing TagRFP-T to generate pHR-ERK^KTR^-TagRFP-T. Murine GRB2 cDNA (kindly provided by Mark Davis) was amplified and subcloned into a pHR plasmid containing TagBFP to generate pHR-GRB2-BFP.

### Cell Line Generation and stimulation

Cell lines were generated via lentiviral transduction. Lentivirus was produced by co-transfecting vectors encoding lentiviral packaging proteins (pMD2.G and p8.91) along with a pHR vector containing a gene of interest into Hek-293T cells plated in 6-well plates (Thermo Fisher Scientific) grown to ∼70% confluence. Transit-IT-293 (Mirrus Bio) was used for all transfections. Viral supernatants were harvested 2 days post-transfection, filtered through a 0.45μm syringe filter (Millex), and concentrated ∼40 fold using Lenti-X Concentrator (Takara Bio Inc.). Viral supernatants were either used immediately, stored at 4 °C for up to one week, or stored at −80 °C for long-term storage. 0.5×10^6^ Ramos cells resuspended in 250 μL 10G-RPMI were mixed at a 1:1 ratio with concentrated supernatant and incubated overnight. Following incubation, viral supernatants were removed by centrifugation, and transduced cells were cultured in 10G-RPMI for one week to recover. Transgene expressing cells were isolated by FACS. Clonal RBD^220^-eGFP/ERK^KTR^-Tag-RFP-T co-expressing cells were isolated by limiting dilution. IgM-deficient Ramos cells were isolated by iterative rounds of sorting Ramos cells negative for cell surface IgM expression (assayed by cell surface staining with APC-conjugated anti-IgM antibody, Biolegend #314509).

Cells were stimulated as indicated in the text and Figure legends. αIgM (Biolegend # 314502), PDBu (CST #12808S) and Concanavalin A (Sigma # C5275) were used at the indicated concentrations and were diluted in imaging media. Methyl α-D-mannopyranoside (αMM) (Sigma #M6882) was used at final concentration of 100mM and a fresh stock diluted fresh in imaging media for every experiment (to guard against contamination).

### Microscopy and Image Analysis

Cells were plated in a well of a glass bottom 96-well glass plate (corning). Wells were coated with 1 mg/mL poly-L-lysine (Sigma) for ∼30 mins, and washed 3X with PBS prior to cell seeding (at a concentration of 0.5 million cells/mL). Images were acquired on a Nikon Eclipse Ti inverted microscope equipped with a Yokogawa CSU-X1 spinning disk confocal, a 60x 1.4 Plan Fluor objective (Nikon) and a Prime95B cMOS camera (Photometics). 405nm, 488nm, 561nm and 640nm (LMM5, Spectral Applied Research) laser lines were used for excitation. CellProfiler (Broad Institute) and Fiji (National Institutes of Health) were used for cell segmentation and image analysis. Excel (Microsoft), Pandas (Wes McKinney), PRISM 7 (GraphPad), and Seaborn (Michael Waskom) were used for data wrangling and visualization. For experiments in cells expressing ERK-KTR, nuclei were labeled with nucblue (Thermo Fisher Scientific) per manufacturer instructions.

Raw microscopy images were background subtracted and corrected for stage drift prior to cell segmentation (to identify cells) and erosion (to remove peripheral pixels and identify the cytoplasm). Membrane intensity of signaling activity reporters (Ras, GRB2) was calculated by taking the inverse of the cytoplasmic intensity. Singe cell intensity traces were normalized by dividing traces by a cell’s integrated mean fluorescence prior to stimulation.

The cytoplasmic-to-nuclear ratio of KTR channel intensity was used to quantify KTR activity. The fluorescence intensity of KTR reporter was quantified in the nucleus (identified by nucblue signal) and the cytoplasm (identified by subtracting nuclear region of the cell from the cell mask). Due to the scarcity of cytoplasmic pixels in Ramos cells, the consistency of cell masking from frame to frame was evaluated and adjusted manually for KTR quantitation. As nuclei occupy a large fraction of the cytoplasmic area in Ramos cells, for experiments in which both Ras reporter and KTR signal was measured, cells were imaged at both a coverslip proximal focal plane (to exclude the nucleus, and improve Ras reporter quantitation, **Figure S1**) and equatorial focal plane (to include the nucleus and facilitate KTR quantitation).

All imaging experiments were performed at least two (and generally three) times. Dead cells were identified by eye and excluded from analysis.

### Statistical Analysis

Two-tailed students’ t-tests were performed using PRISM 7 software (GraphPad). Tests with a *p* value <0.05 were considered statistically significant. In Figures, ns indicates not significant, while * represents p < 0.05, ** represent p<0.01 and *** represents p<0.001. Hartigans’ Dip Test was performed in R (diptest package, Martin Maechler). Tests with a p value <0.05 indicate significant bimodality, while tests with p values >0.05 but 0.1 indicated bimodality with marginal significance. The dip statistic (D) indicates the point in a population distribution in which the maximal difference between peaks in the distribution is achieved. For our purposes, it highlights a plausible separation between populations of cells responding to stimulation.

### Western Blots and Fluorescence activated cell sorting (FACS)

For western blot experiments, cell lines were serum starved for 0.5-1 hour in imaging media at concentration of 0.5 × 10^6^ cells/mL. Following stimulation, cells were immediately transferred to ice, pelleted via centrifugation and resuspended in 200μL ice-cold lysis buffer (1% Triton X-100, 50mM HEPES, pH 7.4, 150mM NaCl, 1.5mM MgCl_2_, 1mM EGTA, 100mM NaF, 10mM Na pyrophosphate, 1mM Na_3_VO_4_, 10% glycerol, freshly supplemented with protease and phosphatase inhibitor tablets (Roche Applied Sciences). Lysates were incubated on ice for 10 minutes and centrifuged at 15,000g for 10 minutes. Protein concentration was measured by BCA Assay (Thermo Fisher Scientific), after which samples were solubilized in 4X Sample Buffer (Bio-Rad Laboratories) and boiled at 100°C for 10 minutes. Sample were either immediately loaded onto gels for Western blot analysis or flash frozen. Bolt 17-well precast 4-12% Bis-Tris Plus gels (Thermo Fisher Scientific) were run at 200V for 25 minutes to resolve proteins. Proteins were transferred from gels to nitrocellulose membranes using a Trans-Blot semi-dry transfer apparatus (Bio-Rad) run at 20V for 1 hour. Membranes were blocked in Odyssey Blocking buffer (Li-Cor) for 1 hour, exposed to primary antibodies diluted according to manufacturer instructions in Odyssey blocking buffer for 4°C overnight, washed in TBST, and incubated in IRDye (Li-Cor) anti-rabbit and anti-mouse secondary antibodies for 30 minutes at room temperature. Following a final wash in TBST, membranes were imaged using a Li-Cor imaging station.

Cell surface staining experiments (IgM, CD22) was performed in FACs buffer (PBS, 5% FBS, 1mM EDTA, 0.1% NAN_3_) for 30 mins at 4 °C. Cells were washed 3X in ice-cold FACs buffer, fixed at room temperature in Cytofix (BD Biosciences), washed three additional times in FACs buffer, and run immediately on a BD LSR II (BD Biosciences). For FACs staining of ERK phosphorylation (ppERK), cells were serum-starved as above in 96 well plates (Corning). A 2X stock of stimulus (IαGM, PDBu, etc…) was used to stimulate cells for the appropriate time. Cells were then fixed at room temperature with Cytofix (BD Biosciences), pelleted by centrifugation (500G for 5 mins), and resuspended in ice-cold methanol to permeabilize cell membranes overnight. Cells were washed 3X in ice-cold FACS buffer and stained with anti-ppERK antibody (CST. #4370) diluted 1:1000 in FACs buffer for 1 hour at room temperature. Cells were washed 3X in FACs buffer and stained with 405 conjugated anti-rabbit secondary antibodies (Abcam # ab175672) for 30 mins. Cells were washed 3 additional times in ice-cold FACS buffer and run immediately on BD LSR II (BD Biosciences). Analysis of flow cytometry data was performed in FlowJo (FlowJo LLC).

## Acknowledgements

We thank Doug Tischer, Kevin Thurley, Mable Lam and members of the Weiner lab for helpful discussions and critical reading of the manuscript. We thank Doug Tischer for cloning the pHR-GRB2-TagBFP construct. This work was supported by NIH grants GM109899 and GM118167 (O.D.W.) and the Center for Cellular Construction (DBI-1548297), an NSF Science and Technology Center.

---

**Figure S1.**
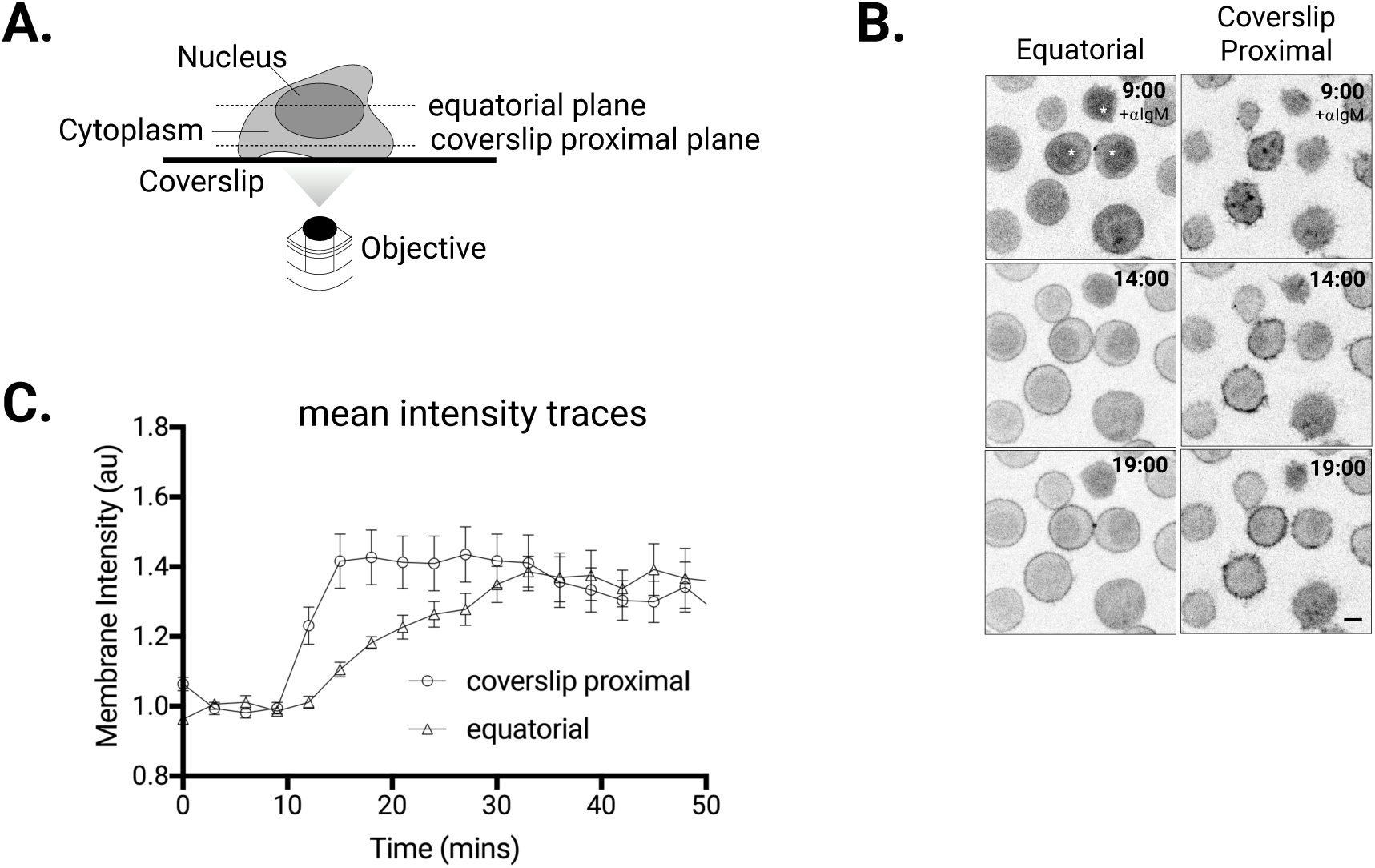
Quantitating Ras signaling dynamics in live cells. **(A)** Schematic depicting imaging set up in which cells are imaged in two focal planes: 1. The equatorial plane, which often includes both the nucleus and cytoplasm, and 2. The coverslip proximal plane, which includes the cytoplasm and generally excludes the nucleus. **(B)** Representative (n>5) time lapse images of RBD^220^-eGFP Ramos cells stimulated with 10μg/mL αIgM (at t = 9 mins). Left column depicts the equatorial focal plane. Nuclei in top left panel are indicated with an asterisk (white). Right column depicts the coverslip proximal focal plane. (C) Mean intensity traces of cells (n=20 cells per focal plane) stimulated as in (B).

**Figure S2.**
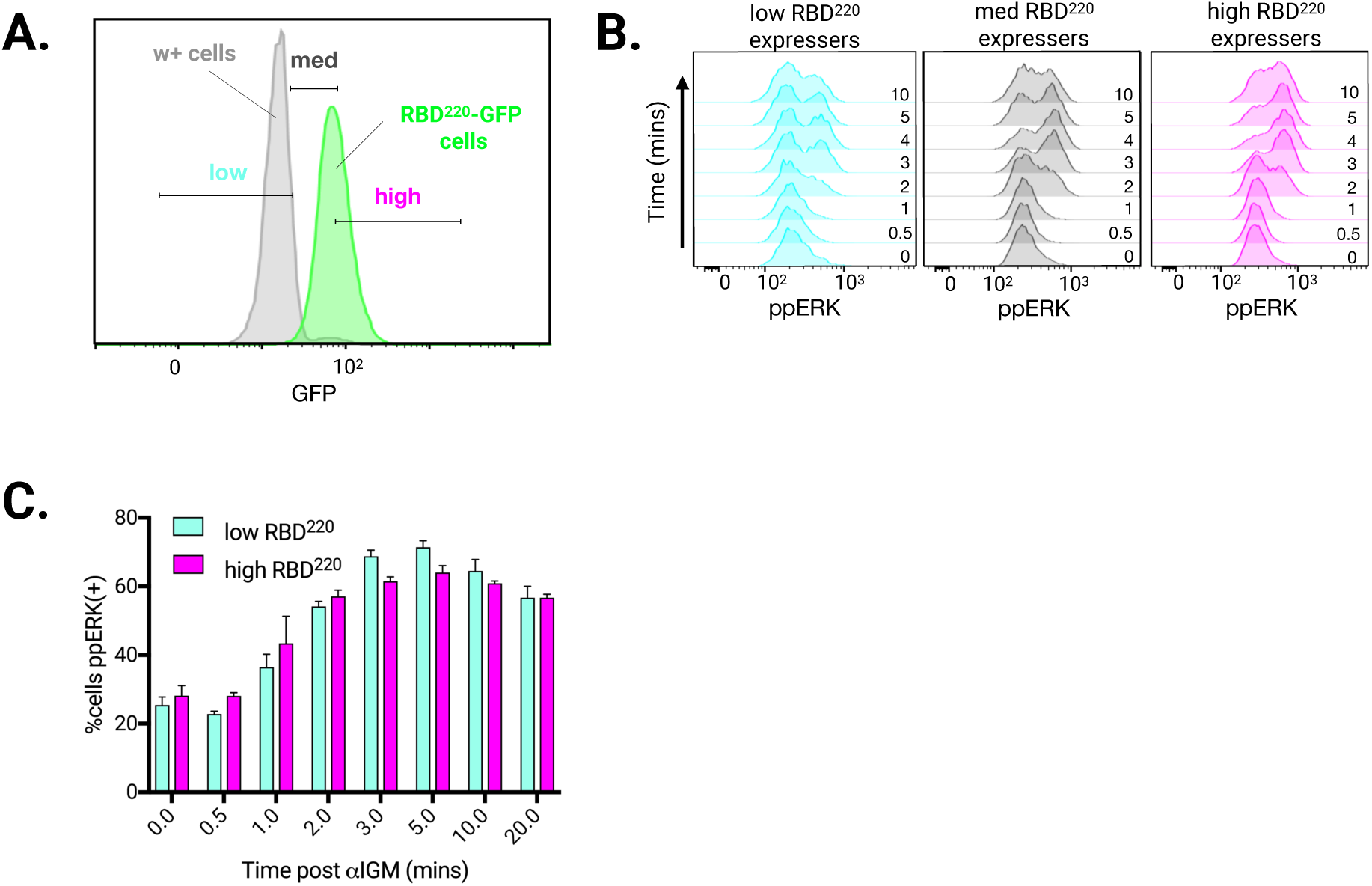
Similar ppERK responses across a wide RBD^220^ expression range in a stable cell line. **(A)** GFP signal in wildtype and RBD^220^-eGFP Ramos cells (100,000 cells each) binned according to GFP signal. **(B)** ERK phosphorylation (ppERK) time course from RBD^220^-eGFP Ramos cells binned based on GFP expression level as in (A), stimulated with 10μg/mL αIgM (at t=0 mins). **(C)** Fraction of RBD^220^ low and RBD^220^ high cells staining positive for ppERK from experiment detailed in (B). Data were pooled from two independent experiments.

**Figure S3.**
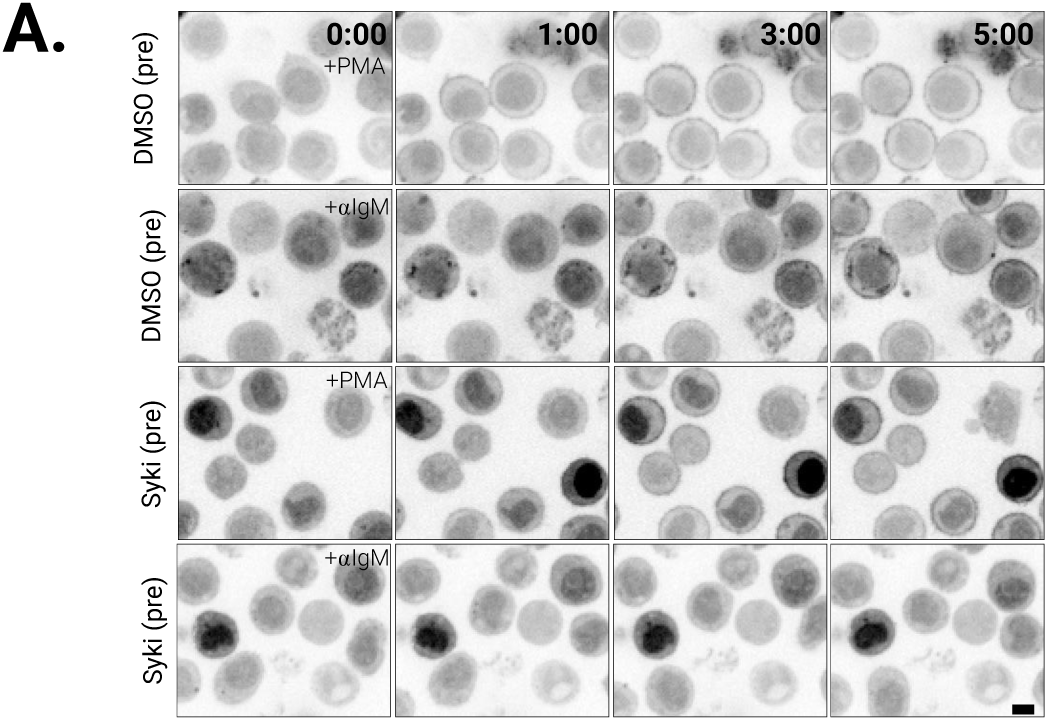
Syk activity is required for αIgM but not PMA-induced RBD^220^ membrane translocation. **(A)** Representative time course images of RBD^220^-eGFP Ramos cells pretreated with either 0.1% DMSO (top two rows) or the Syk inhibitor BAY-61-3606 (10nM) and then stimulated with 10μM PMA (1^st^, 3^rd^ rows) or 10μg/mL αIgM (2^nd^, 4^th^ rows) at t=0mins. Scale bar (black, bottom right panel) is 5μm.

**Figure S4.**
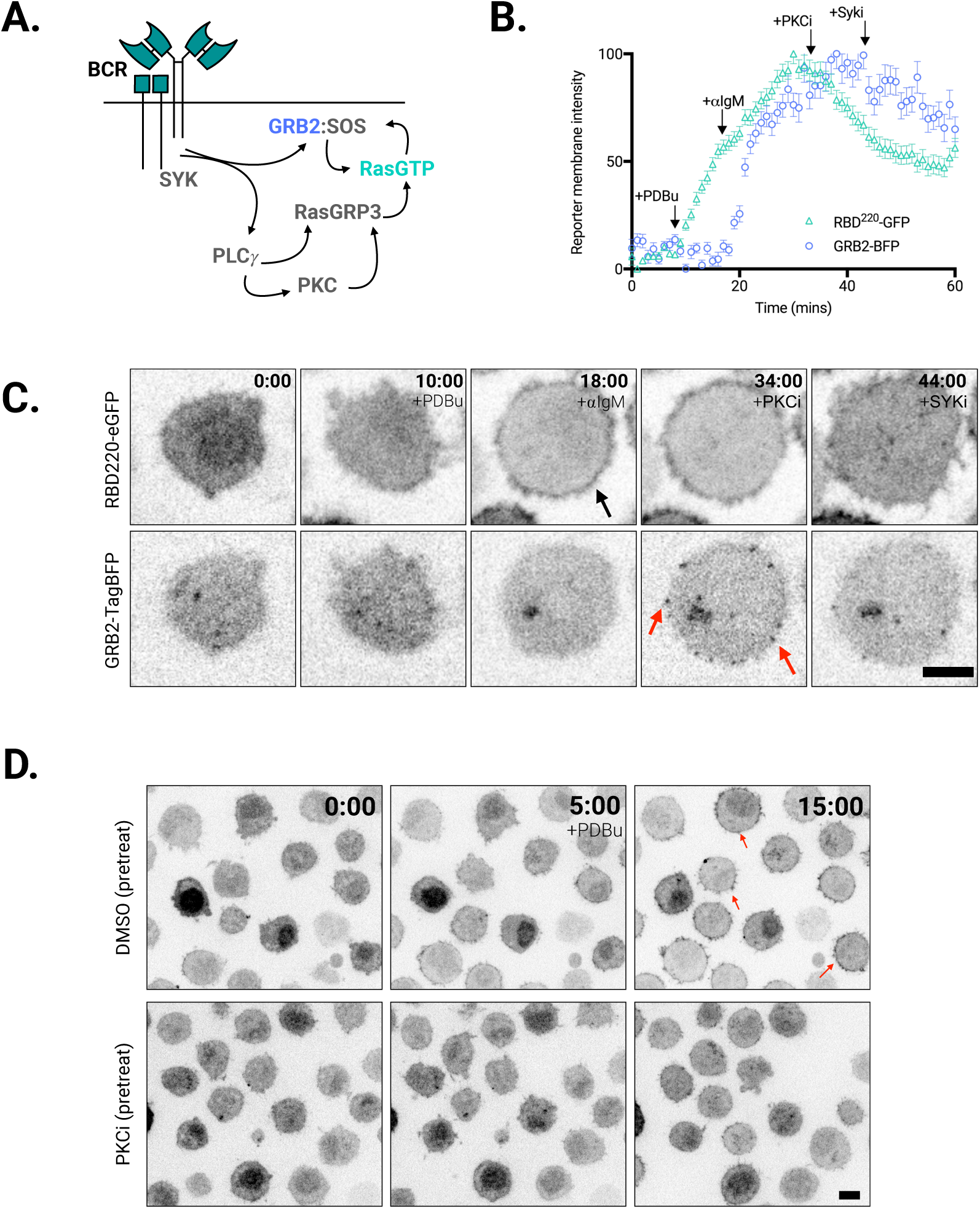
PDBu does not induce GRB2 membrane localization and requires PKC activity for inducing Ras-ERK signaling. **(A)** Schematic depicting BCR triggered Ras activation. While αIgM stimulates both the RasGRP and SOS arm of the pathway, PDBu mimics the PLCγproduct, DAG, thereby solely triggering the RasGRP arm. **(B)** Mean (n=50 cells) time course of GRB2-BFP (blue circles) and RBD^220^-eGFP (green triangles) co-expressing cells stimulated sequentially with 10μM PDBu (at t=10mins), 10μg/mL αIgM (t=18mins), 50nM Gö 6938 (PKCi, at t=34 mins), and 10nM BAY-61-3606 (Syki, at t=44mins). Traces are representative of two independent experiments. Error bars are SEM. **(C)** Single cell stills from cells stimulated as in (B). Arrowheads indicate RBD^220^-eGFP membrane associate (black) and formation of peripheral GRB2-BFP clusters (red) following PDBu and αIgM stimulation, respectively. **(D)** Time course analysis of RBD220-eGFP Ramos cells pre-incubated in 0.1% DMSO (top row) or 50nM Gö 6938 (PKCi bottom row), subsequently treated with 10μM PDBu (t = 5 mins). Arrowheads (red) highlight RBD^220^ membrane translocation.

**Figure S5.**
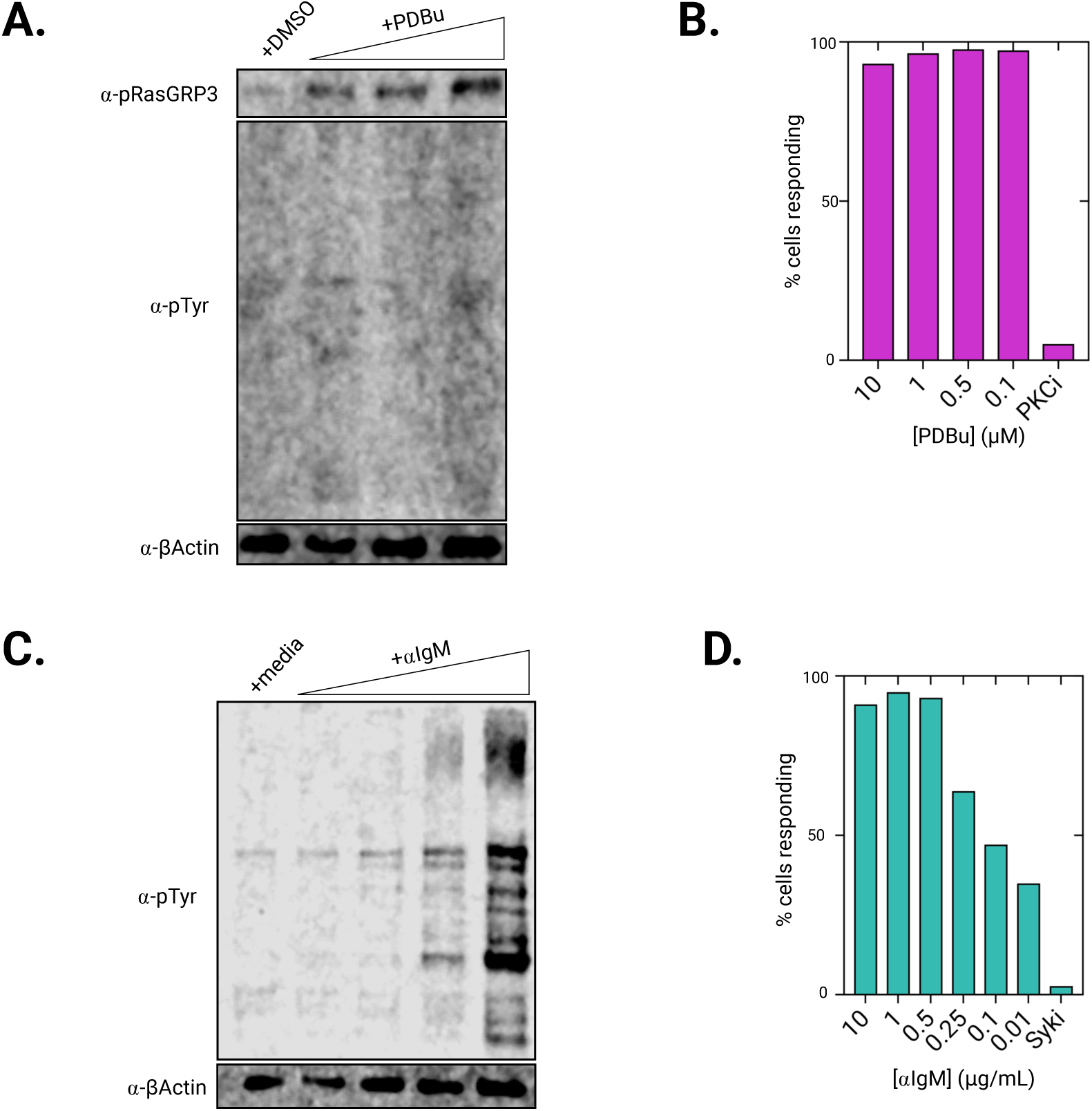
Tuning RasGRP and BCR signaling output by titrating PDBu and αIgM. **(A)** Western blot analysis of RasGRP phosphorylation (T133, a PKC site), global phosphotyrosine (pTyr) and β-Actin (loading control) in wildtype Ramos cells stimulated with DMSO (0.1%) or an ascending concentration of PDBu (0.1μM, 1 μM and 10 μM, left to right). **(B)** RBD^220^-eGFP Ramos cells stimulated with the indicated concentration of PDBu or 50nM Gö 6938 (PKCi). Bars depict the fraction of cells responding. **(C)** Western blot analysis of RasGRP phosphorylation (T133, a PKC site), global phosphotyrosine (pTyr) and β-Actin (loading control) in wildtype Ramos cells stimulated with imaging media or an ascending concentration of αIgM (0.01 μg/mL, 0.1 μg/mL, 1 μg/mL and 10 μg/mL, left to right). (D) RBD^220^-eGFP Ramos cells stimulated with the indicated concentration of αIgM or 10nM Bay-61-3606 (Syki). Bars depict the fraction of cells responding.

**Figure S6.**
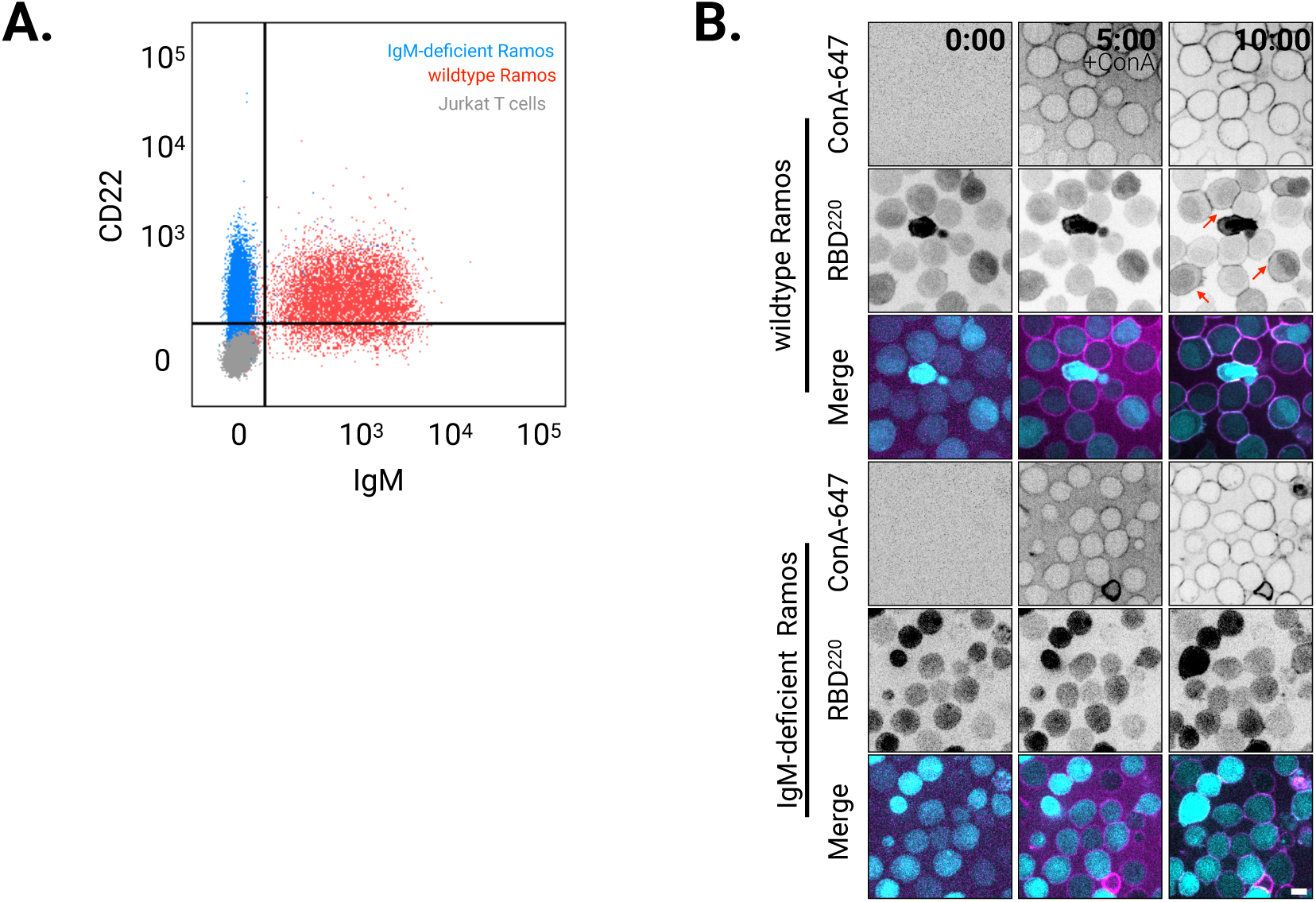
Cell surface BCR expression is required for ConA-induced Ras activation. **(A)** Cell surface staining profiles of IgM and CD22 in Jurkat T cells (grey, negative control), wildtype Ramos cells (red), and IgM-deficient Ramos cells (blue). (**B**) Time course stills of RBD^220^-eGFP expressing wildtype (top three rows) and IgM-deficient (bottom three rows) Ramos cells stimulated with 50μg/mL of 647-labeled ConA (at t=5 mins). Arrows (red) indicate RBD^220^ membrane association. Scale bar (white, bottom right panel) is 5μm.

**Figure S7.**
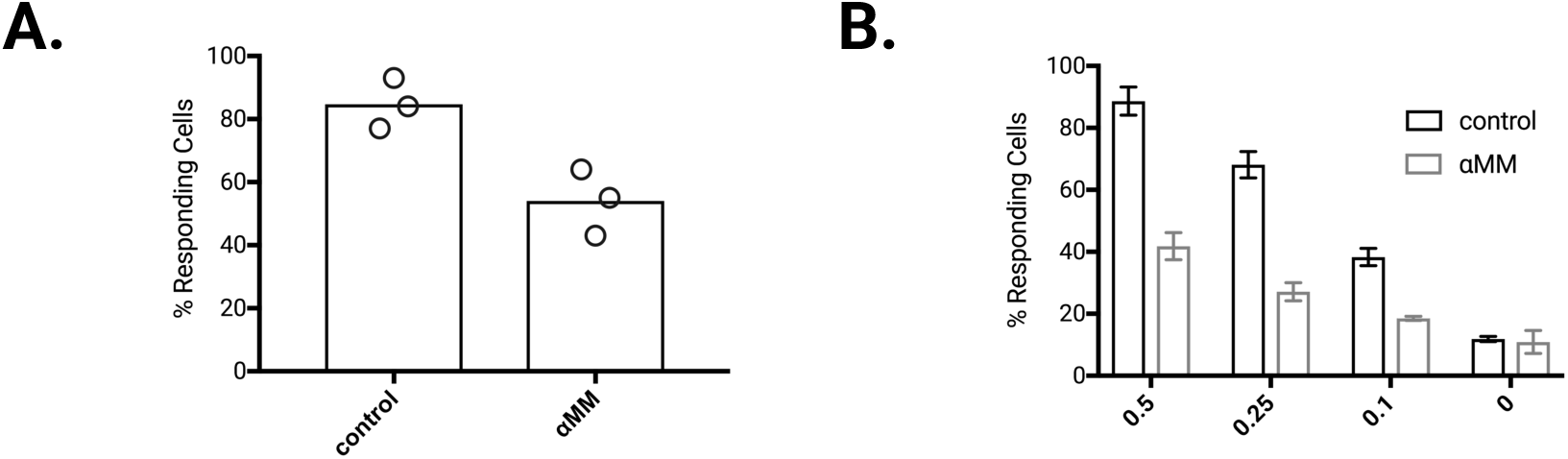
Pretreating cells with αMM partially attenuates αIgM-stimulated Ras responses. **(A)** RBD^220^-GFP expressing Ramos cells were pretreated with either imaging media (control) or 100mM αMM (for 20 mins) prior to stimulation with 10 μg/mL αIgM. The fraction of cells with membrane localized reporter 10 mins after αIgM stimulation is quantified. Each circle represents an independent experiment. At least 20 cells are counted per experiment. (**B)** The fraction of RBD^220^-GFP expressing Ramos cells responding to the indicated doses of αIgM stimulus following pretreatment with either imaging media (control) or 100mM αMM (for 20 mins) is plotted. Error bars are SEM (n=2 independent experiments). 0.25 μg/mL was selected as the minimal stimulus eliciting an RBD^220^-GFP in cells pretreated with αMM for subsequent experiments (figure 4).

**Figure S8.**
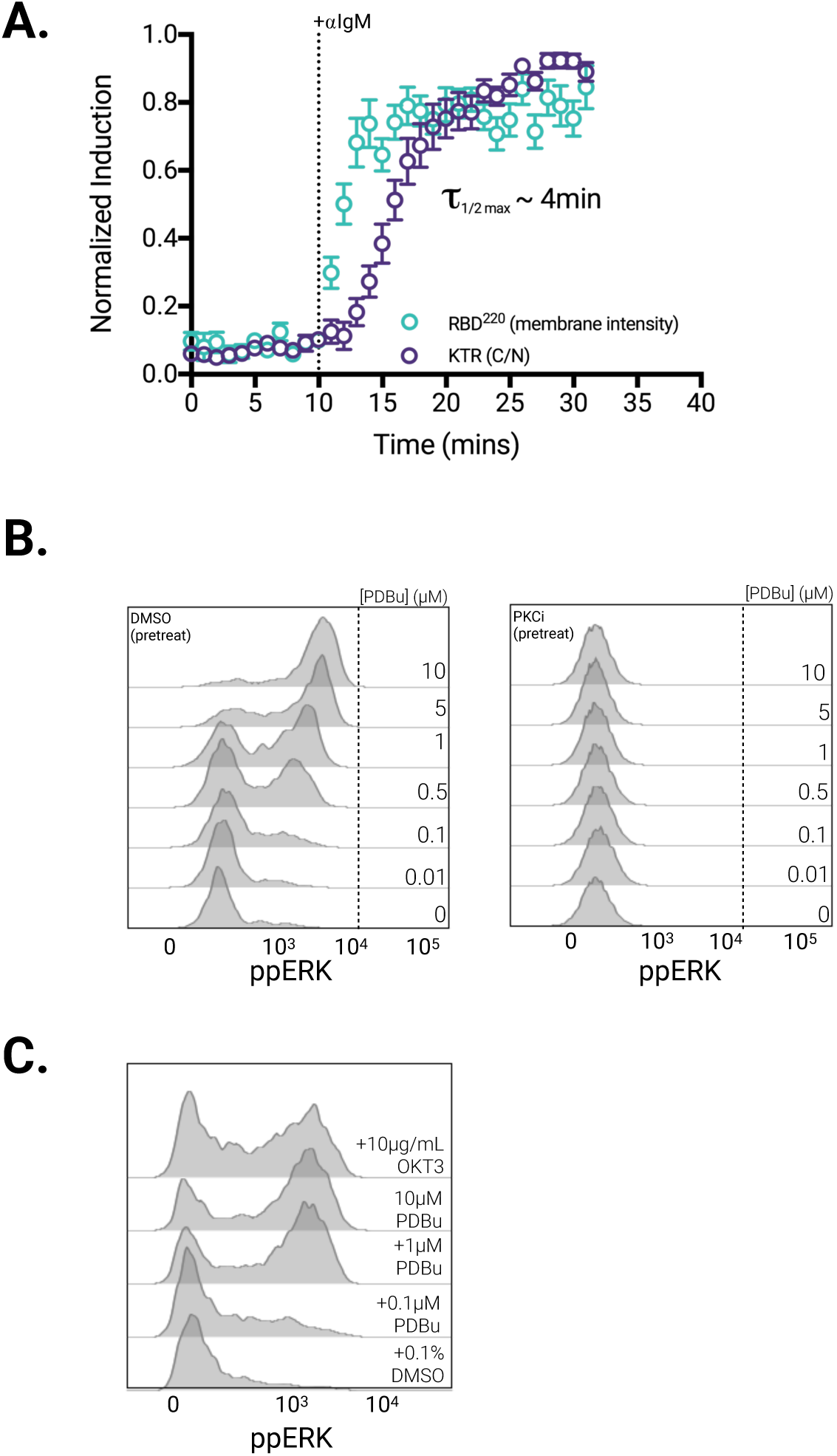
Live and fixed-cell analysis of ERK activation. **(A)** Time-course analysis of RBD^220^ membrane association and ERK-KTR activity from cells co-expressing RBD^220^-eGFP and ERK-KTR-TagRFP-T treated with 10μg/mL αIgM (t = 10mins). The lag time (τ) between half maximal Ras and ERK activity is indicated. **(B)** ERK phosphorylation analysis of wildtype Ramos cells pretreated with DMSO (0.1%, left) or 50nM Gö 6938 (PKCi, right), and treated with the indicated doses of PDBu for 5 mins prior to fixation. Staining from 5000 cells displayed per histogram. **(C)** ERK phosphorylation analysis of wildtype Jurkat T cells treated with the indicated concentrations of DMSO, PDBu or T-Cell receptor cross-linking antibody, OKT3, for 5 mins prior to fixation. Staining from 5000 cells displayed per histogram.

## References

1. Anderson, D. J. et al. (2011) ‘Live-cell microscopy reveals small molecule inhibitor effects on MAPK pathway dynamics’, PLoS ONE, 6(8). doi:10.1371/journal.pone.0022607.

2. Augsten, M. et al. (2006) ‘Live-cell imaging of endogenous Ras-GTP illustrates predominant Ras activation at the plasma membrane’, EMBO Reports, 7(1), pp. 46–51. doi:10.1038/sj.embor.7400560.

3. Bondeva, T. et al. (2001) ‘Structural Determinants of Ras-Raf Interaction Analyzed in Live Cells’, Molecular Biology of the Cell, 13(February), pp. 2323–2333. doi:10.1091/mbc. E02.

4. Boykevisch, S. et al. (2006) ‘Regulation of Ras Signaling Dynamics by Sos-Mediated Positive Feedback’, Current Biology, 16(21), pp. 2173–2179. doi:10.1016/j.cub.2006.09.033.

5. Burger, J. et al. (2009) ‘stimulation High-Level Expression of the T Cell Chemokines CCL3 and CCL4 by Chronic Lymphocytic Leukemia B cells in nurselike cell co-cultures and after BCR stimulation, Leukemia, 113(13), pp. 3050–3059. doi:10.1182/blood-2008-07-170415.

6. Chiu, V. K. et al. (2002) ‘Ras signalling on the endoplasmic reticulum and the Golgi’, Nature Cell Biology, 4(5), pp. 343–350. doi:10.1038/ncb783.

7. Christensen, S. M. et al. (2016) ‘One-way membrane trafficking of SOS in receptor-triggered Ras activation’, Nature Structural and Molecular Biology. Nature Publishing Group, 23(9), pp. 838–846. doi:10.1038/nsmb.3275.

8. Das, J. et al. (2009) ‘Digital Signaling and Hysteresis Characterize Ras Activation in Lymphoid Cells’, Cell. Elsevier Inc., 136(2), pp. 337–351. doi:10.1016/j.cell.2008.11.051.

9. Hartigan, J. A. and Hartigan, P. M. (1985) ‘The Dip Test of Unimodality’, The Annals of Statistics, 13(1), pp. 70–84. doi:10.1214/aos/1176346577.

10. Hibino, K. et al. (2009) ‘A RasGTP-induced conformational change in C-RAF is essential for accurate molecular recognition’, Biophysical Journal. Biophysical Society, 97(5), pp. 1277–1287. doi:10.1016/j.bpj.2009.05.048.

11. Huang, J. et al. (2013) ‘A Single peptide-major histocompatibility complex ligand triggers digital cytokine secretion in CD4+T Cells’, Immunity. Elsevier Inc., 39(5), pp. 846–857. doi:10.1016/j.immuni.2013.08.036.

12. Iversen, L. et al. (2014) ‘Ras activation by SOS: Allosteric regulation by altered fluctuation dynamics’, 345(6192). doi:10.1126/science.1250373.

13. Iwig, J. S. et al. (2013) ‘Structural analysis of autoinhibition in the Ras-specific exchange factor RasGRP1’, eLife, 2013(2), pp. 1–28. doi:10.7554/eLife.00813.

14. Jun, J. E. et al. (2013) ‘Activation of Extracellular Signal-Regulated Kinase but Not of p38 Mitogen-Activated Protein Kinase Pathways in Lymphocytes Requires Allosteric Activation of SOS’, Molecular and Cellular Biology, 33(12), pp. 2470–2484. doi:10.1128/MCB.01593-12.

15. Lorenzo, P. S. et al. (2000) ‘The guanine nucleotide exchange factor RasGRP is a high - affinity target for diacylglycerol and phorbol esters.’, Molecular pharmacology, 57(5), pp. 840–6. doi:10.1124/mol.61.4.759.

16. Malleshaiah, M. K. et al. (2010) ‘The scaffold protein Ste5 directly controls a switch-like mating decision in yeast’, Nature. Nature Publishing Group, 465(7294), pp. 101–105. doi:10.1038/nature08946.

17. Margarit, S. M. et al. (2003) ‘Structural evidence for feedback activation by RasGTP of the Ras-specific nucleotide exchange factor SOS’, Cell, 112(5), pp. 685–695. doi:10.1016/S0092-8674(03)00149-1.

18. Mochizuki, N. et al. (2001) ‘Spatio-temporal images of growth-factor-induced activation of Ras and Rap1’, Nature, 411(6841), pp. 1065–1068. doi:10.1038/35082594.

19. Ogura, Y. et al. (2018) ‘A Switch-like Activation Relay of EGFR-ERK Signaling Regulates a Wave of Cellular Contractility for Epithelial Invagination’, Developmental Cell. Elsevier Inc., 46(2), p. 162–172.e5. doi:10.1016/j.devcel.2018.06.004.

20. Oliveira, A. F. and Yasuda, R. (2013) ‘An Improved Ras Sensor for Highly Sensitive and Quantitative FRET-FLIM Imaging’, PLoS ONE, 8(1), pp. 1–5. doi:10.1371/journal.pone.0052874.

21. Regot, S. et al. (2014) ‘High-sensitivity measurements of multiple kinase activities in live single cells’, Cell. Elsevier Inc., 157(7), pp. 1724–1734. doi:10.1016/j.cell.2014.04.039.

22. Richards, J. D. et al. (2001) ‘Inhibition of the MEK/ERK Signaling Pathway Blocks a Subset of B Cell Responses to Antigen’, The Journal of Immunology, 166(6), pp. 3855–3864. doi:10.4049/jimmunol.166.6.3855.

23. Rubio, I. et al. (2010) ‘TCR-Induced Activation of Ras Proceeds at the Plasma Membrane and Requires Palmitoylation of N-Ras’, The Journal of Immunology, 185(6), pp. 3536–3543. doi:10.4049/jimmunol.1000334.

24. Spencer, S. L. and Sorger, P. K. (2011) ‘Measuring and modeling apoptosis in single cells’, Cell. Elsevier Inc., 144(6), pp. 926–939. doi:10.1016/j.cell.2011.03.002.

25. Teixeira, C. et al. (2003) ‘Integration of DAG signaling systems mediated by PKC-dependent phosphorylation of RasGRP3’, Blood, 102(4), pp. 1414–1420. doi:10.1182/blood-2002-11-3621.

26. Thapar, R., Williams, J. G. and Campbell, S. L. (2004) ‘NMR characterization of full-length farnesylated and non-farnesylated H-Ras and its implications for Raf activation’, Journal of Molecular Biology, 343(5), pp. 1391–1408. doi:10.1016/j.jmb.2004.08.106.

27. Toettcher, J. E., Weiner, O. D. and Lim, W. A. (2013) ‘Using optogenetics to interrogate the dynamic control of signal transmission by the Ras/Erk module’, Cell. Elsevier Inc., 155(6), pp. 1422–1434. doi:10.1016/j.cell.2013.11.004.

28. Weiss, a et al. (1987) ‘Ligand-receptor interactions required for commitment to the activation of the interleukin 2 gene.’, Journal of immunology, 138(7), pp. 2169–2176.

29. Williams, J. G. et al. (2000) ‘Elucidation of binding determinants and functional consequences of Ras/Raf-cysteine-rich domain interactions’, Journal of Biological Chemistry, 275(29), pp. 22172–22179. doi:10.1074/jbc.M000397200.

30. Wilson, M. Z. et al. (2017) ‘Tracing Information Flow from Erk to Target Gene Induction Reveals Mechanisms of Dynamic and Combinatorial Control’, Molecular Cell. Elsevier Inc., 67(5), p. 757–769.e5. doi:10.1016/j.molcel.2017.07.016.

31. Xiong, W. and Jr, J. E. F. (2003) ‘A positive-feedback-based bistable “memory module” that governs a cell fate decision’, 426(November), pp. 460–466. doi:10.1038/nature02119.1.

32. Zheng, Y. et al. (2005) ‘Phosphorylation of RasGRP3 on threonine 133 provides a mechanistic link between PKC and Ras signaling systems in B cells’, Blood, 105(9), pp. 3648–3654. doi:10.1182/blood-2004-10-3916.

33. Zikherman, J. and Au-Yeung, B. (2015) ‘The role of T cell receptor signaling thresholds in guiding T cell fate decisions’, Current Opinion

